# Twin-arginine translocase component TatB performs folding quality control via a general chaperone activity

**DOI:** 10.1101/2020.05.11.089458

**Authors:** May N. Taw, Jason T. Boock, Daniel Kim, Mark A. Rocco, Dujduan Waraho-Zhmayev, Matthew P. DeLisa

**Author notes:** Address correspondence to: Matthew P. DeLisa, Robert Frederick Smith School of Chemical and Biomolecular Engineering, Cornell University, Ithaca, NY 14853.

## Abstract

The twin-arginine translocation (Tat) pathway involves an inbuilt quality control (QC) system that synchronizes proofreading of substrate protein folding with lipid bilayer transport. However, the molecular details of this QC mechanism remain poorly understood. Here, we hypothesized that the conformational state of Tat substrates is directly sensed by the TatB component of the bacterial Tat translocase. In support of this hypothesis, several TatB variants in which the cytoplasmic membrane-extrinsic domain was either truncated or mutated in the vicinity of a conserved, highly flexible α-helical domain were observed to form functional translocases *in vivo* that had compromised QC activity as evidenced by the uncharacteristic export of several misfolded protein substrates. *In vitro* folding experiments revealed that the membrane-extrinsic domain of TatB possessed general chaperone activity, transiently binding to highly structured, partially unfolded intermediates of a model protein, citrate synthase, thereby preventing its irreversible aggregation and stabilizing the active species. Collectively, these results suggest that the Tat translocase may use chaperone-like client recognition to monitor the conformational status of its substrates.

## Introduction

A major challenge faced by all cells is the transport of proteins across tightly sealed, energy-transducing membranes. In prokaryotes, proteins are trafficked across the cytoplasmic membrane using two primary routes: (1) the general secretory (Sec) pathway that requires proteins to be maintained in an unstructured state; and (2) the twin-arginine translocation (Tat) pathway that transports proteins that have already achieved a folded conformation (for reviews, see refs. (1) and (2)). This latter feat is accomplished by a translocase comprised of the integral membrane proteins TatA, TatB, and TatC. TatB and TatC form a receptor complex that binds substrate proteins bearing an N-terminal signal peptide containing a characteristic RR motif (3–5). Substrate binding triggers polymerization of TatA, creating the protein-conducting complex (6, 7). TatA assemblies are believed to form either a ‘pore’ corresponding to the size of the substrate protein (8) or a ‘patch’ that weakens or disorders the membrane bilayer to facilitate protein passage across the membrane (9). The latter model is currently favored as it explains how the translocase accomplishes the difficult task of transporting both small (∼20 Å) and large (∼70 Å) protein structures without opening large holes that would dissipate the proton-motive force (PMF).

Among the many substrates that natively transit the TatABC translocase are cofactor-containing redox proteins (10, 11) and oligomeric complexes (12, 13), all of which involve cytoplasmic assembly prior to export. For these and many other Tat-targeted proteins, proper folding in the cytoplasm with any cofactors in place appears to be a prerequisite for translocation. Indeed, incorrectly folded substrates, including those with even small alterations of substrate conformation, are blocked for export and the rejected molecules are rapidly degraded (14–20). Collectively, these studies point to the existence of a folding quality-control (QC) mechanism that distinguishes the extent of folding and conformational flexibility of Tat substrate proteins, preventing the futile export of those that are misfolded or misassembled. In one notable example, *Escherichia coli* alkaline phosphatase (PhoA) modified with a functional Tat signal peptide was only exported by the TatABC translocase in mutant strains that permitted oxidative protein folding in the cytoplasm and thus generated properly folded PhoA moieties prior to export (16). While not translocated, reduced and misfolded PhoA was still able to specifically associate with the TatBC receptor as revealed by site-specific cross-linking (21–23). However, the binding to TatBC differed between folded and unfolded PhoA precursors, suggesting that the ability to discriminate the folding status of exported proteins may reside within one of these translocase components.

At present, however, very little is known about this QC mechanism or how components of the Tat machinery ‘sense’ the folding state of a protein substrate. Efforts to address this question have leveraged synthetic substrates to help define the conformational cues that are perceived by the translocase and to what extent these features correlate with productive transport. For example, Tat export was hardly detectable when varying-length repeats derived from the FG domain of the yeast nuclear pore protein Nsp1p, which adopts a natively unfolded conformation (24), were heterologously expressed in the presence of native levels of *tatABC* (25). Moreover, when six residues from the hydrophobic core of a globular protein were inserted into each of these repeat constructs, translocation was completely blocked, suggesting that the Tat system proofreads proteins based on surface hydrophobicity. A separate study investigated Tat export of a heterologous single-chain variable (scFv) antibody fragment and mutagenized derivatives with altered surface properties but intact tertiary structures, and found that export efficiency increased with greater structural rigidity (26). Unlike with the FG repeats, the Tat machinery was tolerant of significant changes in hydrophobicity as well as charge on the scFv surface. Based on these results, it was concluded that conformational flexibility of the substrate was the critical attribute discerned by the QC mechanism. The Tat system’s preference for more rigid structures has similarly been demonstrated using *de novo*-designed protein substrates with well characterized differences in the extent of folding. For example, the α_3_ family of designed three-helix-bundle proteins that represent a continuum of folded structures ranging from aggregation-prone (α_3_A) and monomeric molten globules (α_3_B) to well-ordered three-helical bundles (α_3_C and α_3_D) were found to exhibit clear differences in translocation, with increasingly well-folded proteins exported with greater efficiency (27). Nearly identical results were obtained with a panel of four-helix bundle maquette proteins having different conformational flexibility due to the extent of heme *b* cofactor loading (28) and with a collection of Alzheimer’s Aβ42 peptides that were progressively stabilized by point mutations (29).

Using the α_3_ proteins as reporters for a genetic selection, we demonstrated that QC could be inactivated through the isolation of suppressor mutations in the TatABC components, leading to the export of misfolded protein structures that are normally rejected by the wild-type (wt) translocase (27). These findings clearly established that substrate proofreading was, at least in part, executed at the level of the Tat translocase and occurred independently of protein translocation. Strikingly, 21 of the 23 individual QC suppressor (QCS) mutations in TatB were enriched in the membrane-extrinsic portion of TatB (residues 22-171) following the transmembrane helix (Fig. 1a). At present, however, the functional role of TatB and in particular the membrane-extrinsic domain of TatB remain poorly defined with only a handful of studies providing any clues. It has been shown that truncation from the C‐terminus of TatB to form a protein corresponding to just the first ∼50 amino acids still allowed export of different physiological substrates (30, 31) and did not have any measurable effect on TatBC complex formation or stability (31). While contacts between TatC and the folded substrate domain have not been detected, multiple sites in the N-terminal transmembrane and adjacent helical domains of TatB have been identified that contact the major part of a Tat substrate’s surface (3, 23), leading to the proposal of a cage-like structure of the cytosolic TatB domain that transiently accommodates the folded Tat substrate prior to its translocation (23).

**Figure 1.**
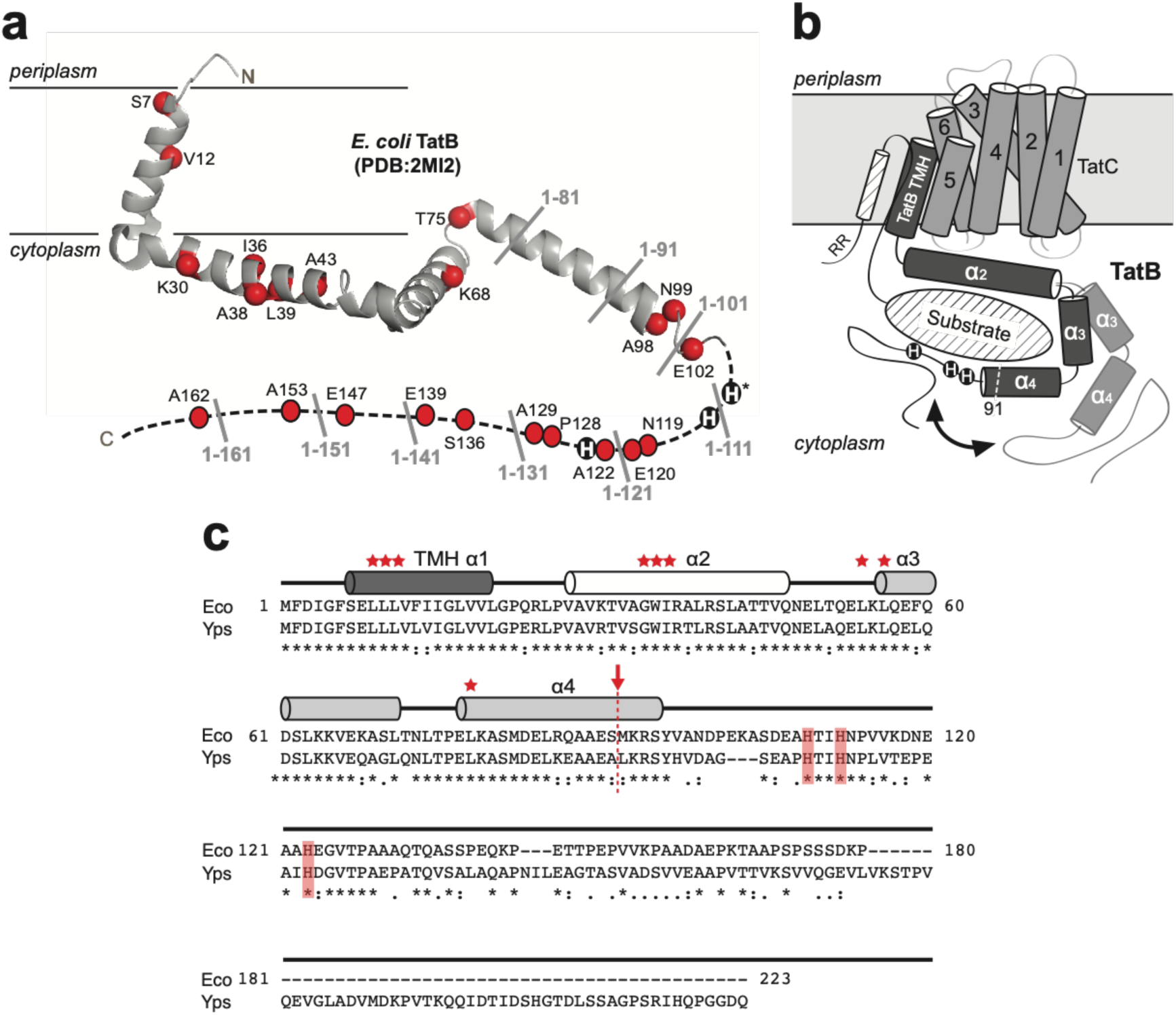
Structure of TatB translocase component. (a) Ribbon diagram of the solution structure of TatB^1-101^ adapted from Zhang *et al*. (32). QCS mutations isolated by Rocco *et al*. (27) are marked as red balls (except for H109 which is marked with asterisk). Locations of truncation sites in 10-residue increments from the C-terminus are labeled in gray. Histidines at positions 109, 112, and 123 are shown as black circles. (b) Model of possible TatB-substrate interactions adapted from Ulfig *et al*. (35). The framed light gray box represents the lipid bilayer, while a TatC monomer is depicted by six transmembrane helices, and a TatB monomer is depicted with a membrane-embedded transmembrane helix (TMH α1), a strongly amphipathic helix (α2), and two highly hydrophilic and flexible helices (α3 and α4). The cytosolic α3 and α4 helices encapsulate a folded substrate protein (diagonal hatched oval), with possible movement of the helices depicted by black arrow and dashed line cylinders. It should also be pointed out that the TatBC complex functions as a receptor for the N-terminal signal peptide, which is defined by a twin-arginine motif (RR) and h-region α-helix (diagonal hatched cylinder). For clarity, the cartoon does not account for the well-known signal peptide-TatBC interaction. Also for clarity, only one TatB monomer (dark gray) and one TatC monomer (light gray) are shown, but the model could easily accommodate higher order oligomeric structures such as the tetrameric TatBC complex described by Lee *et al*. (36). One possible scenario for QC could be that upon binding of the signal peptide and N-terminal part of the substrate, the C-terminal domain of TatB dynamically wraps around the substrate and performs conformational proofreading. (c) Multiple sequence alignment of TatB proteins from γ-proteobacteria *E. coli* and *Y. pseudotuberculosis* generated using CLUSTALW. Asterisks indicate identical amino acids, colons indicate conservation between amino acids with strongly similar properties, and periods indicate conservation between amino acids with weakly similar properties. The α-helical regions were adapted from the solution structure of TatB (32) and are represented as cylinders. The truncation point after residue 91 is marked with red arrow/dashed line and the histidines at positions 109, 112, and 123 are shaded red. Residues shown by Maurer *et al*. (23) to form contacts with folded substrates are marked with red stars.

Based on these earlier findings, we hypothesized here that the cytoplasmic domain of TatB performs a QC function that operates at a distinct stage of the transport cycle and that may involve chaperone-like client recognition to monitor the conformational status of its substrates. To test this hypothesis, we assessed the consequences of incrementally truncating residues from the C-terminus of TatB to remove membrane-extrinsic portions of the protein. This analysis uncovered greatly shortened TatB variants that assembled into functional translocases with the uncharacteristic ability to export misfolded protein substrates, indicating QC activity had been compromised. Using a set of *in vitro* assays for testing chaperone function, we discovered that the entire membrane-extrinsic domain of TatB interacted with highly structured, partially unfolded intermediates but not unstructured, completely unfolded intermediates of a classical chaperone substrate protein, citrate synthase (CS). Taken together, our results provide a possible explanation for how the TatB component of the translocase might sense flexible motions and conformations of the substrate, which could then be transmitted to other components of the Tat machinery for preventing transport across the membrane.

## Results

### Folding quality control is dependent on the membrane-extrinsic domain of TatB

The *E. coli* TatB protein is 171 amino acids in length and adopts an extended “L-shape” conformation consisting of four α-helices: a transmembrane helix (TMH) α1 (residues 7-20); an amphipathic helix (APH) α2 (residues 27-47); and two hydrophilic helices α3 (residues 56-71) and α4 (residues 77-96) (Fig. 1a and c) (32). Whereas the TMH and APH segments are relatively rigid, these latter two helices display notably higher mobility, which may allow TatB to bind substrate proteins with different sizes and shapes (Fig. 1b). The C-terminal region of the protein from residue 96 onwards is predicted to have a predominantly random coil conformation. To test our hypothesis that the cytoplasmic membrane-extrinsic domain of TatB following the TMH is involved in folding QC, we performed truncation analysis by removing up to 140 residues from the C-terminus of TatB in 10-residue increments (Fig. 1a) and evaluating the resulting mutants using a genetic assay that directly links Tat folding QC activity with antibiotic resistance (27). This assay involves a panel of fusion constructs comprised of: (i) an N-terminal Tat signal peptide derived from trimethylamine *N-*oxide reductase (spTorA); (ii) one of the designed three-helix-bundle proteins (α_3_A, α_3_B, α_3_C, and α_3_D; Fig. 2a) that exhibit progressively greater conformational rigidity (33, 34); and (iii) C-terminal TEM-1 β-lactamase (Bla).

**Figure 2.**
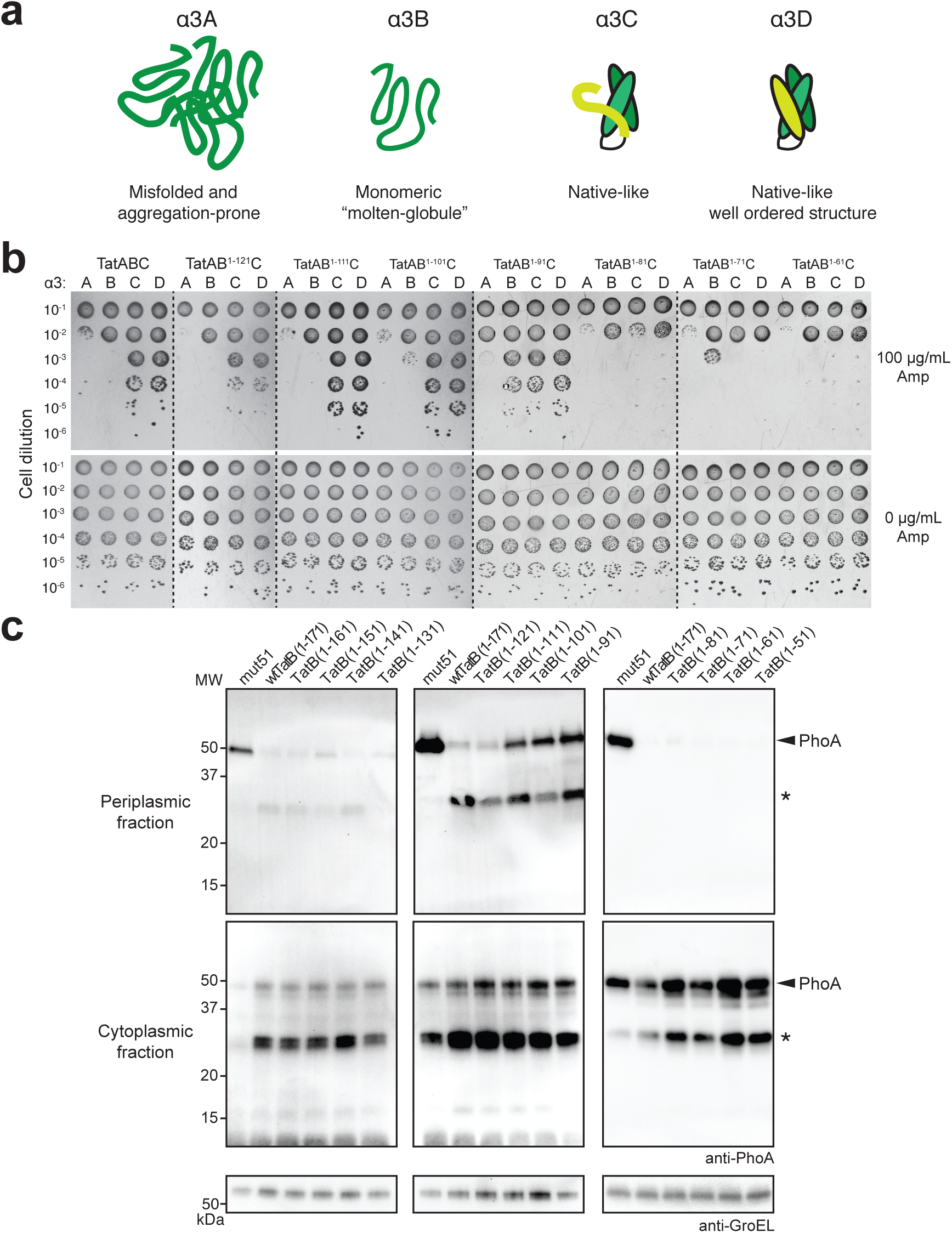
Truncation of membrane-extrinsic domain abrogates QC activity. (a) Schematic representation of the α3 family of designed three-helix-bundle proteins developed by DeGrado and coworkers (33, 34) which represent a continuum of folded structures, ranging from aggregation-prone (α3A) and monomeric molten globular (α3B) to native-like, well-ordered (α3C and α3D). (b) Resistance of serially diluted DADE cells co-expressing spTorA-α_3_-Bla chimeras (A, B, C or D) along with a Tat operon plasmid encoding either wt TatB or one of the TatB variants. Cells were spotted on LB-agar plates containing either 100 μg/mL Amp or 25 μg/mL chloramphenicol (Cam; 0 μg/mL Amp). Dashed lines denote different LB-agar plates that were all generated and imaged at the same time. (c) Western blot analysis of cytoplasmic and periplasmic fractions prepared from DADE cells co-expressing Tat-targeted PhoA from pTorA-AP along with a Tat operon plasmid encoding either wt, one of the TatB variants, or mut51. An equivalent number of cells was loaded in each lane. PhoA was probed with anti-PhoA antibody while anti-GroEL antibody confirmed equivalent loading in each lane. Asterisk indicates degraded spTorA-PhoA.

The ampicillin (Amp) resistance conferred by these different constructs to a *tat*-deficient *E. coli* strain called DADE (MC4100 Δ*tatABCD*Δ*Ε*) (56) carrying a plasmid-encoded copy of the wt TatABC operon was in the following order (from highest to lowest): α3D ≈ α3C >> α3B > α3A (Fig. 2b), which was in close agreement with the resistance conferred by these constructs to the isogenic parental strain MC4100 that expressed TatABC natively (27). Nearly identical resistance profiles were observed for DADE cells expressing TatB proteins lacking as many as 50 C‐terminal amino acids (TatB^1-121^) (Fig. 2b), indicating that most of the random coil portion of TatB was dispensable for both QC and translocase activities. Removal of 60 or 70 C-terminal amino acids from TatB (TatB^1-111^ or TatB^1-101^, respectively, which each comprise the TMH, the adjacent α-helical domains, and the first 5-15 residues of the random coil region) resulted in a very low but detectable level of increased growth for α3B but not α3A relative to growth conferred by translocases comprised of wt TatB (Fig. 2b). These mutants also exhibited slightly increased export of α3C and α3D. More remarkably, when TatB was C-terminally truncated by 80 amino acids (TatB^1-91^, which is disrupted in solvent exposed helix α4), even stronger QC suppression was observed with greatly increased export of α3B and modestly increased export of α3A (Fig. 2b), reminiscent of the phenotype that was previously ascribed to class II-type QCS translocases (27). Further truncation of TatB resulted in severely diminished export of all spTorA-α3-Bla constructs. Notably, there were no apparent growth defects for any of the strains when plated in the absence of Amp (Fig. 2b).

To determine whether the Amp resistance was attributable to translocases that retained native function, the truncated TatB proteins were assessed for the ability to export two native Tat substrates, namely the *N*-acetylmuramoyl-L-alanine amidases AmiA and AmiC. In *E. coli*, these enzymes cleave the peptide moiety of *N*-acetylmuramic acid in peptidoglycan and contribute to daughter cell separation by helping to split the septal murein (37). Mutations that impair Tat export lead to mislocalization of AmiA and AmiC, rendering *E. coli* sensitive to SDS and disrupting cell division (38). The cell division defect results in the formation of cell chains ranging from 6 to 24 cells in length. Indeed, DADE cells carrying an empty plasmid formed characteristic chains (more than 6 cells per chain) whereas DADE cells carrying a plasmid-encoded copy of the wt TatABC operon showed no visible cell-division defects (**Supplementary Fig. 1a**). Importantly, DADE cells expressing TatB proteins that were C-terminally truncated by as many as 120 amino acids (TatB^1-51^) divided properly (**Supplementary Fig. 1a**), in agreement with previous studies that also reported export of physiological Tat substrates in the presence of significantly truncated TatB proteins (30, 31). Only after removal of 130 or more amino acids from the C-terminus of TatB did we observe cell division defects associated with the absence of TatB (**Supplementary Fig. 1a**). Hence, whereas Tat export was contingent on a TatB protein comprised minimally of the TMH and amphipathic helix α2, QC required a longer TatB that included the highly flexible α3 and α4 helices in addition to the TMH and α2.

To further explore the role of the membrane-extrinsic domain of TatB in folding QC, the truncated TatB proteins were assessed for their ability to export misfolded PhoA. Previously, we demonstrated that PhoA was only exported by the Tat pathway if its native disulfide bonds had formed to generate the correctly folded molecule in the cytoplasm prior to export (16). This outcome required the genetically modified cytoplasm of *E. coli* strain DR473, which has the unnatural capacity to catalyze disulfide bond formation in the cytoplasmic compartment. More recently, we found that reduced, misfolded PhoA could be exported from the normally reducing cytoplasm of wt *E. coli* cells that co-expressed QCS translocases (27). Here, we hypothesized that TatB^1-91^, TatB^1-101^, and TatB^1-111^ would similarly promote export of reduced PhoA given the ability of each to export the molten globular α3B substrate and, in the case of TatB^1-91^, also the aggregation-prone α3A. Indeed, wt cells expressing translocases containing TatB^1-91^, TatB^1-101^, and TatB^1-111^, but not wt TatB or any other truncated TatB variants, were able to export reduced PhoA as revealed by Western blot analysis (Fig. 2c). It should be noted that the amount of PhoA exported by TatB^1-91^, TatB^1-101^, and TatB^1-111^ was visibly lower than that achieved with the previously isolated class I QCS translocase mut51 (Fig. 2c).

### Histidine residues in membrane-extrinsic domain are essential for folding QC

Of the 21 individual QCS mutations that were previously isolated in the membrane‐extrinsic domain of TatB, one of these was a histidine to asparagine substitution at residue 109 (27). This residue and two additional nearby histidines (H112 and H123) formed a histidine ‘patch’ that was located just after the solvent exposed helix α4 and in the vicinity of the truncation sites after residues 91, 101, and 111 that caused relaxation of QC (Fig. 3a). This patch was intriguing in light of similar occurrences of histidine residues in molecular chaperones that are reported to play important roles in substrate binding and release (39–43). To determine the importance of these residues to QC, we performed spot plate analysis of DADE cells expressing hybrid translocases comprised of TatB variants in which these histidines were individually or collectively mutated to alanine. In each case, cells expressing the TatB variants were significantly more resistant to antibiotic in the context of α3B than cells expressing wt TatB (Fig. 3b). Moreover, all of the histidine substitution mutants were able to export misfolded PhoA, with periplasmic accumulation exceeding that observed for TatB^1-91^-containing translocases and rivaling that of the strong QCS translocase mut51 (Fig. 3c). However, these residues were not essential for physiological export as DADE cells expressing hybrid translocases comprised of these TatB variants exported AmiA and AmiC to the periplasm as evidenced by restoration of normal cell division (**Supplementary Fig. 1a**, shown for TatB^H112A^ and the triple mutant). Taken together, these results demonstrate that the clustered histidine residues occurring just after the TMH-adjacent α-helices play an important role in the folding QC mechanism.

**Figure 3.**
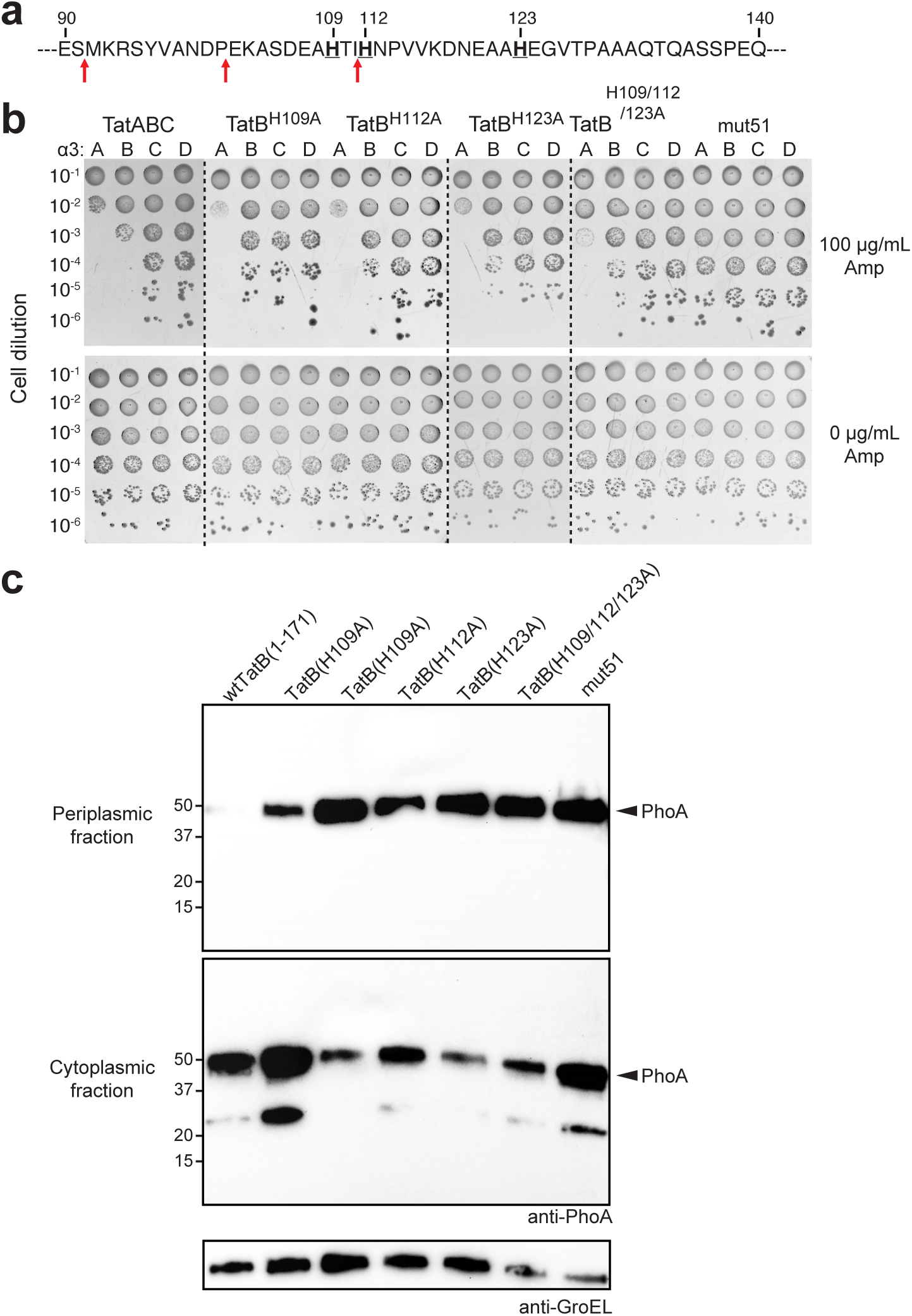
Histidine patch in membrane-extrinsic domain of TatB regulates folding QC. (a) Sequence of residues 90-140 of the membrane-extrinsic domain of TatB. Bold underline font denotes histidines at positions 109, 112, and 123; red arrows indicate QC-relevant truncation sites. (b) Serially diluted DADE cells co-expressing spTorA-α_3_-Bla chimeras (A, B, C or D) along with a Tat operon plasmid encoding wildtype TatB, one of the TatB variants, or the mut51 translocase were spotted on LB-agar plates containing either 100 μg/mL Amp or 25 μg/mL Cam (0 μg/mL Amp). Dashed lines denote different LB-agar plates that were all generated and imaged at the same time. (c) Western blot analysis of cytoplasmic and periplasmic fractions prepared from DADE cells co-expressing Tat-targeted PhoA from pTorA-AP along with either wt TatABC, mut51, or one of the histidine mutants as indicated. An equivalent number of cells was loaded in each lane. PhoA was probed with anti-PhoA antibody while anti-GroEL antibody confirmed equivalent loading in each lane.

### *Yersinia pseudotuberculosis* TatB regulates QC activity in *E. coli*

To determine whether TatB-mediated QC is conserved in other bacteria, we attempted to functionally reconstitute the mechanism in *E. coli* cells using an orthologous TatB from *Y. pseudotuberculosis* (YpTatB) in place of *E. coli* TatB. YpTatB shares ∼57% sequence homology with *E. coli* TatB, most of which occurs in the TMH, amphipathic helix, and the two hydrophilic α-helices. However, starting at residue Y96 in the membrane-extrinsic domain, the YpTatB ortholog diverges significantly and also has a 49-residue C-terminal extension that is absent in *E. coli* TatB (Fig. 1b). Despite these differences, YpTatB was able to form hybrid translocases with *E. coli* TatA and TatC that exported AmiA and AmiC to the periplasm as evidenced by restoration of normal cell division in DADE cells (**Supplementary Fig. 1b**). Encouraged by this result, we next investigated whether wild-type YpTatB could exert QC by regulating export of the Tat-targeted α3 reporter constructs. When spot plated on Amp, DADE cells co-expressing the different α3 constructs along with TatA(YpTatB)C exhibited resistance profiles that were indistinguishable from cells expressing *E. coli* TatABC, with α3A-expressing cells the least resistant and α3D-expressing cells the most (**Supplementary Fig. 2a**). We also observed that heterologous TatA(YpTatB)C translocases were able to efficiently export folded PhoA from the oxidizing cytoplasm of redox-engineered *E. coli* but rejected misfolded PhoA for export from a normal reducing cytoplasm, mirroring the ability of *E. coli* TatABC translocases to regulate export of PhoA in a folding-dependent manner (**Supplementary Fig. 2b**). Taken together, these results confirm that folding QC activity is functionally conserved within the YpTatB ortholog despite the significant divergence of its C-terminal random coil sequence.

To determine whether the membrane-extrinsic domain of YpTatB was similarly important for QC as it was for *E. coli* TatB, we created a truncation variant of YpTatB in which all C-terminal residues after A91 were removed (YpTatB^1-91^). When we expressed YpTatB^1-91^ in DADE, predominantly singlet cells were observed (**Supplementary Fig. 1b**) indicating the formation of hybrid translocases that could export AmiA and AmiC. Similar to its truncated *E. coli* counterpart, YpTatB^1-91^ was able to increase the level of misfolded PhoA that was exported in cells having a reducing cytoplasm (**Supplementary Fig. 2b**), providing further evidence that QC activity is encoded within the membrane-extrinsic portion of TatB.

### Membrane-extrinsic domain of TatB functions *in vitro* as a molecular chaperone

Molecular chaperones are well known to recognize distinct conformational states of their client proteins. We hypothesized that TatB may utilize a similar chaperone-like activity involving its membrane-extrinsic domain to directly monitor the conformational state of substrates. To investigate the potential chaperone functions of this domain, we expressed and purified a truncation mutant of TatB in which the TMH (residues 1-21) was deleted (TatB^22-171^), yielding a soluble protein comprised of the entire cytoplasmic domain of TatB following the TMH. Analysis by SDS-PAGE and gel filtration indicated that TatB^22-171^ was purified from cell lysates to >90% purity and was primarily tetrameric (**Supplementary Fig. 3a and b**), consistent with the known oligomerization state of full-length TatB (23, 36). To investigate the potential chaperone function of purified TatB^22-171^, we employed a set of *in vitro* assays using a classical chaperone substrate protein, mitochondrial citrate synthase (CS), which has served as a standard measure of molecular chaperone activity (44–48). An attractive feature of CS as a model substrate is that distinct unfolding intermediates can be accessed experimentally, which help to shed light on a chaperone’s functional mechanism. We first evaluated the ability to suppress the thermally-induced aggregation of CS. In the absence of molecular chaperones, CS rapidly and irreversibly aggregates at 43°C, a temperature that resembles heat shock *in vivo* (44, 45). Indeed, CS was completely aggregated within 15 min of incubation at 43°C as monitored by light scattering at 500 nm (Fig. 4). An equimolar ratio of purified TatB^22-171^ to CS was sufficient to almost completely prevent thermal aggregation of CS, a level of suppression that was on par with that achieved by equimolar amounts of the molecular chaperones GroEL and Hsp90 (Fig. 4). The stoichiometry of TatB^22-171^’s chaperone action was on par with that of known chaperones (44, 45). To verify that the chaperone activity of TatB was a specific effect, an equimolar amount of bovine serum albumin (BSA) was used as a control but failed to inhibit CS aggregation (Fig. 4). When the TatB^22-171^ construct was truncated to remove 80 C-terminal residues (TatB^22-91^) or when its three histidine residues were mutated to alanine (TatB^22-171[H109/112/123A]^), the ability to prevent CS aggregation was greatly diminished (Fig. 4), consistent with the inactivation of QC observed *in vivo* for these TatB variants. Using the soluble cytoplasmic domain of YpTatB (YpTatB^22-220^), which purified as an apparent trimer (**Supplementary Fig. 3a and b**), we observed concentration-dependent prevention of CS aggregation that was largely abolished when this domain was truncated after residue A91 (Fig. 4), akin to the results obtained with *E. coli* TatB.

**Figure 4.**
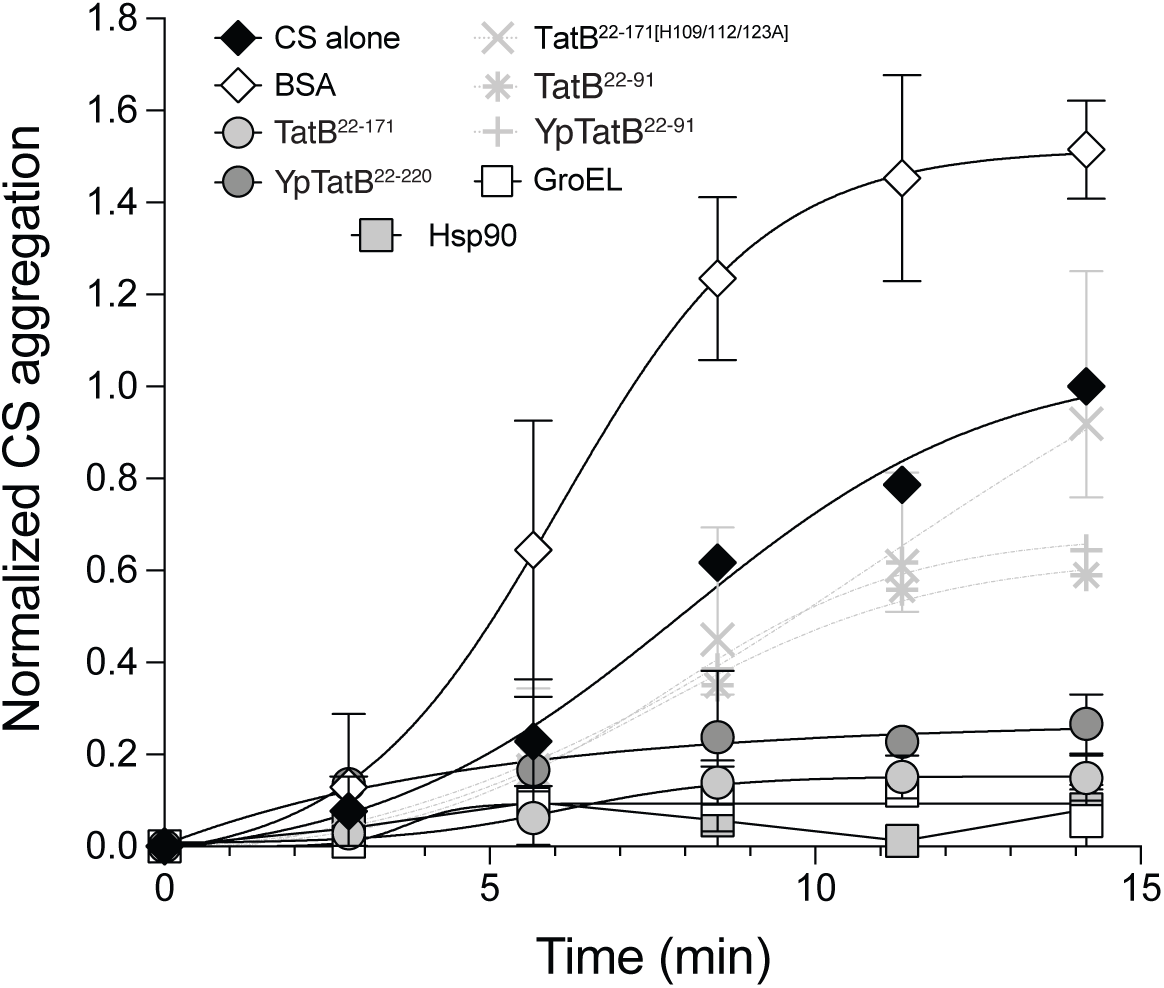
Membrane-extrinsic domain of TatB suppresses thermal aggregation of CS. CS was diluted to a final concentration of 0.15 μM into prewarmed 40 mM HEPES-KOH, pH 7.5, at 43°C in the absence (CS alone) or presence of 0.15 μM of the following proteins: TatB^22-171^, TatB^22-91^, TatB^22-171[H109/112/123A]^, YpTatB^22-220^, YpTatB^22-91^, *E. coli* GroEL or yeast Hsp90 as indicated. To exclude unspecific protein effects, control experiments in the presence of 0.15 μM bovine serum albumin (BSA) were conducted. Light scattering measurements were performed by measuring absorbance at 500 nm. Data are the average of biological replicates and the error bars represent the standard deviation of the mean.

To further characterize the ability of TatB^22-171^ to productively interact with highly structured but partially unfolded intermediates of CS (so-called early unfolding intermediates (47)), we studied its effect on the thermal inactivation of CS. Following incubation at 43°C, the enzymatic activity of CS alone or in the presence of an 8-fold molar excess of the non-chaperone BSA was decreased to less than 10% of its initial value in 7 min and was almost completely inactivated by 15 min (Fig. 5a and b). Consistent with previous findings (47), the addition of 1 mM oxaloacetate, a substrate of CS, or an equimolar amount of the molecular chaperone Hsp90 from yeast exerted a stabilizing effect (Fig. 5a). Likewise, both TatB^22-171^ and YpTatB^22-220^ significantly slowed CS inactivation in a concentration-dependent manner, with an 8-fold molar excess of these proteins resulting in a ∼7-11-fold increase in the half-time of inactivation (Fig. 5b and c). Even equimolar amounts of these proteins were sufficient to measurably slow down the inactivation process. Importantly, neither TatB^22-91^ or YpTatB^22-91^ exhibited a stabilizing effect (Fig. 5b and c), further highlighting the importance of the complete set of TMH-adjacent α-helices in the chaperone-like behavior of TatB. It should also be noted that the triple mutant also exhibited impaired thermo-protection albeit not to the same extent as the C-terminally truncated variants (Fig. 5b).

**Figure 5.**
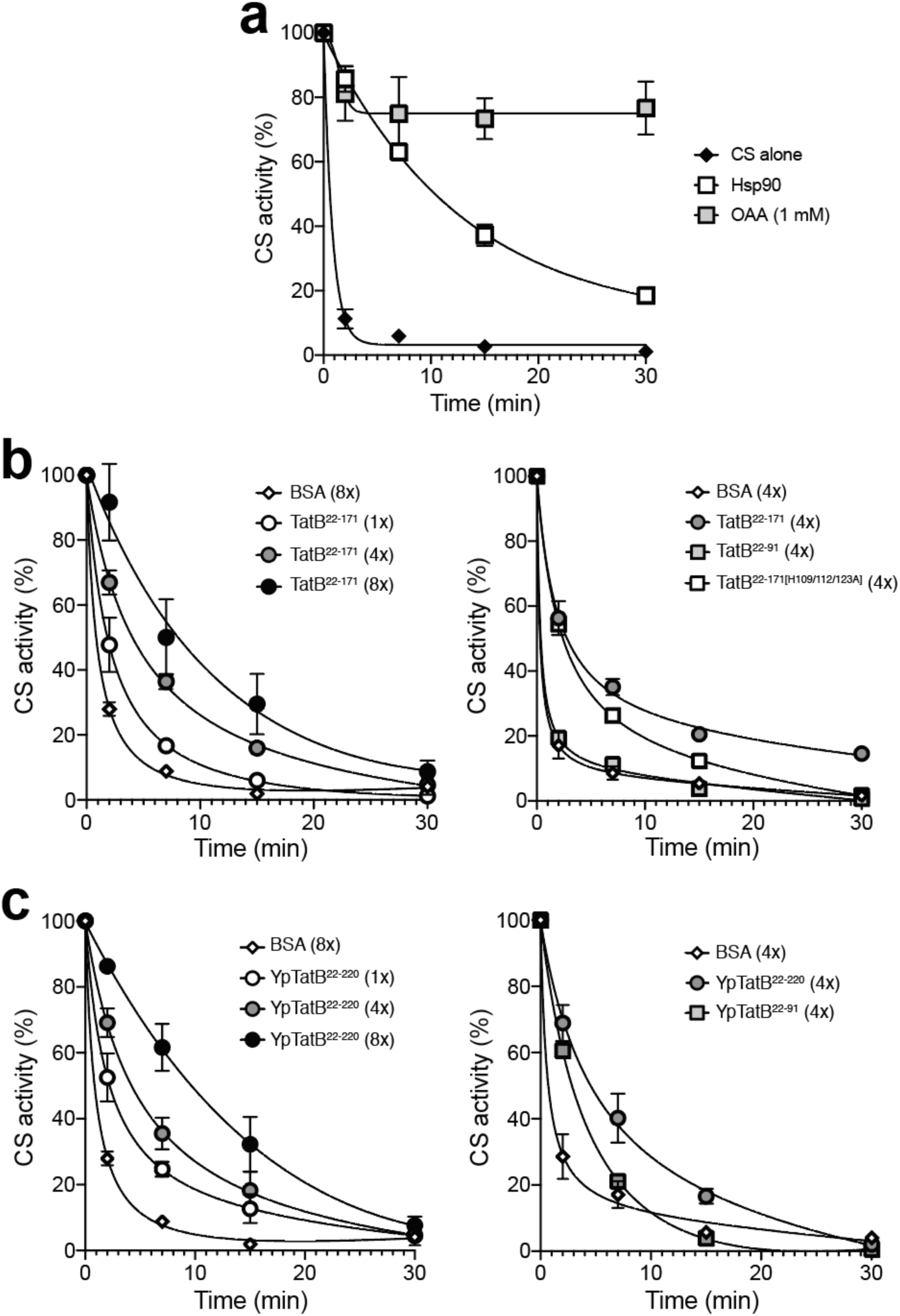
Influence of membrane-extrinsic domain of TatB on thermal inactivation of CS. (a) CS (0.15 μM) was incubated at 43°C either alone or in the presence of 1 mM oxaloacetate (OAA) or equimolar yeast Hsp90 (0.15 μM). (b) Same as in (a) but in the presence of TatB^22-171^ (1x: 0.15 μM; 4x: 0.6 μM, 8x: 1.2 μM), TatB^22-91^ (4x: 0.6 μM), or TatB^22-171[H109/112/123A]^ (4x: 0.6 μM). (c) Same as in (b) but with YpTatB^22-220^ and YpTatB^22-91^. BSA at a concentration of 0.6 μM (4x) or 1.2 μM (8x) served as negative controls. At the times indicated, aliquots were withdrawn and the activity was determined as described. The solid lines represent single exponential functions fit to the experimental data using Prism 8 software. Data are the average of biological replicates and the error bars represent the standard deviation of the mean.

We next investigated whether TatB^22-171^ was able to interact with more significantly unfolded intermediates such as chemically or thermally denatured proteins. To this end, we first assayed the ability of TatB^22-171^ to inhibit aggregation of refolding CS following complete denaturation in guanidine hydrochloride. In this assay, CS is diluted from denaturant into refolding buffer, upon which it immediately aggregates as a result of the high local concentration of aggregation-sensitive folding intermediates (46). In our hands, only GroEL but not TatB^22-171^ or YpTatB^22-220^ was able to prevent aggregation of chemically denatured CS (**Supplementary Fig. 4a**). We also determined whether TatB^22-171^ was capable of reactivating heat-denatured CS. For this experiment, CS was first inactivated at 43°C for 30 min and then cooled to ∼23°C, after which CS activity in the presence of equimolar amounts of TatB^22-171^ or control proteins was measured over time. Again, the molecular stabilizer, oxaloacetate, was able to promote CS reactivation whereas TatB^22-171^ had no measurable effect under the conditions tested (**Supplementary Fig. 4b**). The observation that TatB^22-171^ and YpTatB^22-220^ could neither prevent aggregation of chemically denatured CS nor reactivate heat-denatured CS indicated that these TatB proteins do not recognize completely unfolded or otherwise highly unstructured intermediates of CS. Therefore, we conclude that TatB-mediated sensing of substrate foldedness involves preferential interactions with more structured protein conformations, akin to some small heat shock proteins (45).

## Discussion

The conformational state of a protein during membrane translocation limits and defines the possible mechanisms by which it is delivered to its final destination. In the case of the bacterial Tat pathway, it is now firmly established that substrate proteins cannot be transported until the native structure is reached – failure to incorporate cofactors, assemble with a biological partner, or otherwise attain a correctly folded structure is well known to thwart translocation, leading to accumulation of non-exported precursor forms of the substrate in the cytosol that in some cases are removed by proteolytic degradation (14–20). Because the time and energy required to export folded proteins is high, with transit half-times on the order of minutes (14) and energetic costs equivalent to 104 molecules of ATP (49), Tat export must be carefully regulated so that futile export of misfolded or misassembled proteins is avoided. However, while privileged export of properly folded structures by bacterial Tat translocases has long been known, how different folding states of substrate proteins are distinguished and how this information is integrated into the active transport cycle is poorly understood. For some native Tat substrates that assemble complex cofactors (*e.g.*, the molybdenum cofactor-containing enzyme TorA), QC involves substrate-specific chaperones that coordinate cofactor loading with membrane translocation (50, 51) as well as proteases that eliminate immature or misfolded precursors (52). Yet, it remains enigmatic how QC is accomplished for Tat substrates that do not have dedicated folding catalysts or for artificial substrates (*e.g.*, PhoA, the α3 protein family) that do not natively transit the Tat pathway but are nonetheless subjected to stringent QC when targeted to the bacterial Tat translocase (16, 27, 28). Interestingly, in the case of PhoA, crosslinking studies and suppressor genetics suggest that QC is executed by the translocase directly (21, 27), but a detailed explanation of exactly how this is accomplished is lacking.

In the present study, we provide several lines of evidence that the membrane-extrinsic domain of TatB proofreads the conformational state of protein substrates *in vivo*. First, TatB proteins that were truncated at a site within solvent exposed helix α4, or at sites just after α4 in the early part of the random coil domain, assembled into functional translocases; however, these translocases uncharacteristically exported misfolded protein substrates including reduced PhoA, indicating that QC was impaired. Second, a ‘histidine patch’ overlapping with these truncation sites was identified that upon alanine substitution also triggered export of misfolded proteins. While the specific role of these residues remains to be determined, it is worth noting that a handful of molecular chaperones possess Zn^2+^-binding or pH-sensing histidine residues, which can trigger dramatic conformational changes including structural stabilization/destabilization and dimerization and can also modulate chaperone activity (39–43). It is therefore intriguing to ask whether the membrane-extrinsic histidine cluster might play a similar role in TatB. It is also worth noting that whereas the complete ensemble of TMH-adjacent helices was required for QC activity, significantly shorter TatB proteins comprised of just the TMH and amphipathic α2 helix were capable of forming stable receptor complexes with TatC (31) and exporting different physiological substrates as was observed here and elsewhere (30, 31). These findings suggest that QC function is separable from translocation and appears to be encoded in the highly flexible α3/α4 helices and the ∼5-15 downstream residues, although contributions of the TMH and α2 to the QC mechanism cannot be ruled out. The observed “floppiness” of helices α3 and α4 (32), in particular, would allow TatB to sample a large conformational space and facilitate interaction with numerous structurally diverse substrate proteins.

The relaxation of QC caused by these different genetic alterations in the membrane-extrinsic domain led us to hypothesize that TatB may employ chaperone-like client recognition to discriminate the conformational state of substrate proteins. In support of this hypothesis, *in vitro* folding experiments revealed a general chaperone activity associated with the entire membrane-extrinsic domain of TatB, which preferentially interacted with highly structured but partially unfolded intermediates of CS but not more significantly unfolded intermediates that were highly unstructured. This chaperone-like behavior was abrogated for TatB variants truncated within the α4 helix or mutated in the histidine patch, further reinforcing the notion that QC activity depended on the entire set of α-helical domains and downstream histidine residues. Importantly, these *in vitro* results were not only in agreement with our *in vivo* data but also entirely consistent with previous crosslinking studies demonstrating that (i) TatB but not TatC formed molecular contacts with surface-exposed residues of folded and partially folded precursors but not completely unfolded proteins (3, 23); and (ii) interaction sites in TatB occurred at multiple positions along the TMH and all three adjacent α-helices (Fig. 1b) and were detected for two structurally unrelated substrates, namely PhoA and the native Tat substrate SufI (23).

Our discovery of chaperone-like activity in the TatB translocase component is reminiscent of the cytoplasmic ATPase motor protein SecA, which reportedly uses general chaperone activity to execute a QC function for the Sec pathway (53). However, unlike TatB, SecA binds to unfolded nascent polypeptides, delivering signal peptide-bearing proteins to the SecYEG translocase while stimulating the folding of proteins that do not contain a signal peptide so as to exclude them from SecYEG, which requires its substrates to be unfolded (54). It is particularly noteworthy that both pathways employ bifunctional proteins to effectively integrate proofreading of substrate conformation with the important functional roles of binding the signal peptide and early mature part of precursor proteins in the case of TatB and membrane targeting and energetic driving of unfolded protein translocation in the case of SecA. The importance of such coordinated QC operations cannot be understated as the constant equilibrium of folded and unfolded polypeptides in the cytoplasm could create significant problems for the cell if improper substrate conformations were not efficiently excluded from these highly specialized export mechanisms.

The chaperone function of TatB suggests that posttranslational targeting to the translocase at the cytoplasmic membrane is a dynamic process involving binding and release of substrates. Upon functionally targeting the Tat translocase by virtue of an RR-containing signal peptide, Tat precursors are known to become surrounded by multiple TatB proteins in a cage-like fashion (23). This arrangement allows the α-helical domains, in particular the highly mobile α3 and α4 helices, to productively engage the surface of the substrate. Our finding that TatB does not interact with highly unstructured versions of CS along with an earlier report that TatB does not interact with hidden sites in folded Tat precursors including TorA-PhoA (23) clearly indicate that TatB contacts only the surface of folded Tat precursors. It would follow, therefore, that misfolded substrates such as reduced TorA-PhoA (21) that are interrogated by TatB are not completely unfolded. Indeed, the absence of the two essential disulfide bonds in reduced TorA-PhoA has been suggested to cause either a local unfolding or a generally relaxed conformation with an enlarged surface area (23), providing important clues as to the key QC attributes that are sensed by TatB. In light of these results and the notable preference of the Tat system for more rigid substrate structures (26–29), we favor a model for QC in which TatB uses its inbuilt chaperone-like activity to differentially interact with bound substrates as a function of their structural flexibility, with flexible motions of the substrate inducing changes in TatB’s binding affinity and/or conformation. Because TatB interactions occur with both folded and misfolded substrates and do not depend on the PMF (21, 23), which energizes the membrane translocation step, the TatB-mediated QC step must precede Tat-dependent translocation or the clearance of translocation-incompetent substrates. This notion is supported by crosslinking studies in which the proximity between TatB and bound precursor was lost upon transmembrane transport of the folded precursor (23), highlighting the temporary nature of substrate engagement by TatB. These transient interactions likely induce distinct TatB conformations that are transmitted to other components of the Tat machinery, effectively gating the assembly of the oligomeric TatA pore/patch and preventing transport across the membrane. For example, interactions between TatB and an improperly folded substrate may impede the position switching of TatB with TatA, a recently described phenomenon that represents a critical step in driving the assembly of an active Tat translocase and that only occurs in response to translocation-competent substrates (55).

Collectively, our results suggest that Tat QC involves initial encounters between the membrane-extrinsic domain of TatB and the substrate at the membrane surface. These interactions appear to mimic those that occur between molecular chaperones and their clients and requires the entire α-helical portion and early unstructured region of the membrane-extrinsic domain. Importantly, these results provide a possible explanation for how the Tat translocase senses the structural flexibility of its substrates and subsequently uses this information to restrict protein transport across the membrane.

## Materials and Methods

### Bacterial strains and plasmids

All bacterial strains and plasmids used in this study are listed in **Supplementary Table 1**. *E. coli* strain DH5α was used for all molecular biology while strain DADE(DE3) was used for expression and purification of TatB and YpTatB proteins while BL21(DE3) was used for expressing all other recombinant proteins. *E. coli* strain DADE (MC4100 Δ*tatABCD*Δ*tatE*) (56) lacking all of the *tat* genes was used for all Tat translocation experiments, unless otherwise noted. To determine export of PhoA under reducing cytoplasmic conditions, either DHB4 or MCMTA was used. To compare these with PhoA export under an oxidizing cytoplasm, either DR473 or DRB was used.

To generate plasmids encoding the different TatB truncation mutants, the gene encoding *E. coli* TatB was PCR amplified using the same forward primer to amplify the 5’ end of the gene and a reverse primer that successively truncated the sequence in increments of 10 codons at a time from the 3’ end until only the N-terminal 21 residues remained. The resulting PCR products were cloned into pTatABC-XX (27) in place of the gene encoding wt TatB using the XbaI and XhoI restriction sites that flanked the gene. Plasmids pTatA(YpTatB)C and pTatA(YpTatB^1-91^)C were generated identically except that the PCR products inserted into pTatABC-XX were either full-length or truncated copies of the gene encoding TatB from *Y. pseudotuberculosis.* To generate plasmid pMAF10-TatB, the gene encoding *E. coli* TatB was PCR amplified with primers that introduced XbaI and SphI restriction sites at the 5’ and 3’ end of the gene, respectively, and the resulting PCR product was ligated into the same sites in plasmid pMAF10. An identical strategy was used to create plasmid pMAF10-YpTatB for expressing TatB from *Y. pseudotuberculosis*. To construct pET-TatB^22-171^ and pET-TatB^22-91^, full-length or truncated versions of the gene encoding *E. coli* TatB were PCR amplified with primers that introduced NdeI and HindIII restriction sites at the 5’ and 3’ end of the gene, respectively, and the resulting PCR products were ligated into the same sites in plasmid pET-22b(+) (Novagen). An identical strategy was used to generate plasmids pET-YpTatB^22-220^ and pET-YpTatB^22-91^ for expressing and purifying full-length or truncated versions of TatB from *Y. pseudotuberculosis*. Histidine to alanine substitutions in TatB were introduced by site-directed mutagenesis of plasmids pTatABC-XX and pET-TatB^22-171^.

### Selective plating of bacteria

Bacterial plating was performed as described (27, 29). Briefly, DADE cells harboring one of the pTatABC-XX plasmids and one of the pSALect-α3 plasmids (27) were grown overnight at 37°C in LB medium supplemented with 20 μg/mL tetracycline (Tet) and 30 μg/mL chloramphenicol (Cam). The next day, an equivalent number of cells were harvested from each culture (normalized to an Abs_600_ ≈ 1.0), resuspended in fresh LB medium without antibiotics, and subsequently serially diluted by factors of 10 in sterile 96-well plates. Aliquots of 5 μL from each well were spotted onto LB-agar plates containing Tet and Cam (control) or a specific concentration of Amp. After drying, plates were incubated overnight at 30°C.

### Microscopy

Cultures were inoculated in LB with appropriate antibiotics from freshly transformed strains, grown overnight at 37°C, and subcultured the next day for an additional 4-5 h. A wet mount of live bacterial cells was prepared by using approximately 5 μL of culture and applying a coverslip before being imaged under oil immersion using a Carl Zeiss Axioskop 40 optical microscope with a Zeiss 100x/1,30 Oil Plan-NEOLUAR lens and SPOT FLEX digital camera (Diagnostic Instruments).

### Subcellular fractionation and Western blot analysis

Overnight-grown cells were subcultured 50-fold in Luria-Bertani (LB) medium containing antibiotics and allowed to grow at 37°C until absorbance at 600 nm (Abs_600_) of ∼0.5-0.7 was reached, at which time the cultures were induced with 1 mM IPTG and incubated for an additional 2-3 h at 30°C. An equivalent number of cells were harvested from each culture (normalized to an Abs_600_ ≈ 1.0) and centrifuged at 2,000 x g. Subcellular fractionation was performed using the ice-cold-osmotic shock method (16). The pelleted cells were resuspended in 1 mL fractionation buffer consisting of 30 mM Tris-HCl, pH 8.0, 1 mM EDTA, and 0.58 M sucrose and left at room temp for 10 min. Samples were centrifuged at 9,200 x g for 10 min and resuspended in 150 μl ice-cold 5 mM MgSO4 and kept on ice for 10 min. Samples were again centrifuged at 4°C for 10 min at 16,000 x g. The supernatant was collected as the periplasmic fraction. The remaining pellet was washed once in PBS buffer, resuspended in 300 μl BugBuster® (MilliporeSigma) to lyse cells, and centrifuged once more, with the resulting supernatant collected as the cytoplasmic fraction. Periplasmic and cytoplasmic fractions were separated electrophoretically using SDS-PAGE gels after which Western blotting was performed according to standard protocols using the following antibodies: anti-PhoA (Millipore MAB1012 diluted 1:2,500), anti-mouse-HRP (Abcam ab6789 diluted 1:5,000), anti-GroEL (Sigma G6532 diluted 1:30,000), and anti-rabbit-HRP (Abcam ab205718 diluted 1:5,000).

### Protein purification

Plasmids encoding different TatB and YpTatB variants under the control of the T7 promoter were used to freshly transform *E. coli* BL21(DE3) cells. Transformed cells were inoculated into LB with 100 μg/mL Amp and incubated overnight at 37°C. Strains were subcultured 50-fold into fresh LB with Amp and grown to an Abs_600_ ∼0.5-0.8, at which point they were induced with 0.1 mM IPTG and grown overnight at 30°C. Harvested cells were lysed using a homogenizer (Avestin) and then centrifugued at 12,000 x g for 30 min, with the supernatant collected as the soluble fraction. This fraction was incubated with Ni-NTA resin in equilibration buffer (PBS with 10 mM imidazole, pH 7.4) for 1-2 h at 4°C. Proteins were washed and eluted using PBS with 40 mM imidazole and 250 mM imidazole, pH 7.4, respectively. Total protein concentrations were measured using the Bradford assay with bovine serum albumin (BSA) as standard. Size exclusion chromatography (SEC) was performed using an ÄKTA FPLC system with Superdex75 column (GE Healthcare). Fractions were collected and analyzed by electrophoretic separation on SDS-PAGE gels and subsequent staining with Coomassie blue.

### CS aggregation assays

Porcine CS from Sigma (C3260) was dialyzed into 40 mM HEPES-KOH, pH 7.5. CS was further purified as described previously (45), subjected to SEC, and concentrated using a 30-kDa cut-off concentrator and centrifuged at 15,000 x g to remove insoluble products. Porcine CS prepared in this way was utilized for all *in vitro* folding assays. Thermal aggregation of CS was performed as described previously (47). Briefly, buffer consisting of 40 mM HEPES-KOH, pH 7.5 was equilibrated at 43°C with stirring in the presence or absence of different test proteins (purified TatB and YpTatB variants) and control proteins including *E. coli* GroEL (Sigma C7688), yeast Hsp90 (Enzo Life Sciences ALX-201-138-C025), or BSA (Sigma A7030). Light scattering was measured at 500 nm using a PTI spectrofluorometer with thermostatted cell holder. Both emission and excitation slits were set to 2 nm for the first 5 min to ensure that the putative chaperone alone did not result in aggregation and to achieve a baseline measurement. Next, CS was added to a final concentration of 0.15 μm, and light scattering measurements were performed for a further 15 min to track the thermal aggregation.

The aggregation of chemically denatured CS was assayed as described (45). Briefly, CS was denatured in 6 M guanidinium chloride in 50 mM Tris-HCl, pH 8.0 and left at room temperature for 2 h to ensure complete unfolding of CS. Then, buffer consisting of 40 mM HEPES-KOH, pH 7.5 was equilibrated at 25°C with stirring in the presence or absence of different test proteins (purified TatB and YpTatB variants) and control proteins (GroEL or BSA), and light scattering was measured for 5 min to determine the baseline. Light scattering was measured using the same conditions as the thermal aggregation assay with the exception that the temperature was held constant at 25°C. Renaturation of denatured CS was initiated by diluting CS 100-fold to 0.15 μm in each sample and light scattering measurements were continued for a further 6 min.

### CS inactivation and reactivation assays

TatB variants or BSA was added to 40 mM HEPES-KOH, pH 7.5 at varying concentrations. CS then was added to the sample at a final concentration of 0.15 μm and mixed well. Before the sample was heated, an aliquot was taken to denote the initial amount of CS activity. The samples were then heated to 43°C and aliquots were removed at given time points. Activity in these aliquots was measured as described previously (45) with the exception that 40 mM HEPES-KOH, pH 7.5 was used. Activity measurements were performed at 25°C using a microplate reader (Tecan). For CS reactivation, CS inactivation was performed for 30 min at 43°C as described above. Samples containing CS in the presence or absence of chaperone were allowed to cool to room temperature for 5 min before, after which CS activity was measured periodically over the course of 1 h using a microplate reader (Tecan).

## Acknowledgements

We thank Dr. Tracy Palmer for strains and plasmids used in this work. This work was supported by the National Science Foundation (grants # CBET-0449080 and CBET-1605242 to M.P.D.), the National Institutes of Health (grant # CA132223A (to M.P.D.), the New York State Office of Science, Technology and Academic Research Distinguished Faculty Award (to M.P.D.), and a Royal Thai Government Fellowship (to D.W.-Z.).

## Author Contributions

M.N.T. designed research, performed research, analyzed data, and wrote the paper. J.T.B., M.A.R., and D.W-Z. designed research, performed research, and analyzed data. M.P.D. designed and directed research, analyzed data, and wrote the paper.

## Competing Interests Statement

All authors declare no competing interests.

## Notes

### Competing Interest Statement

The authors have declared no competing interest.

