## Supplementary Information for "Twin-arginine translocase component TatB performs folding quality control via a general chaperone activity"

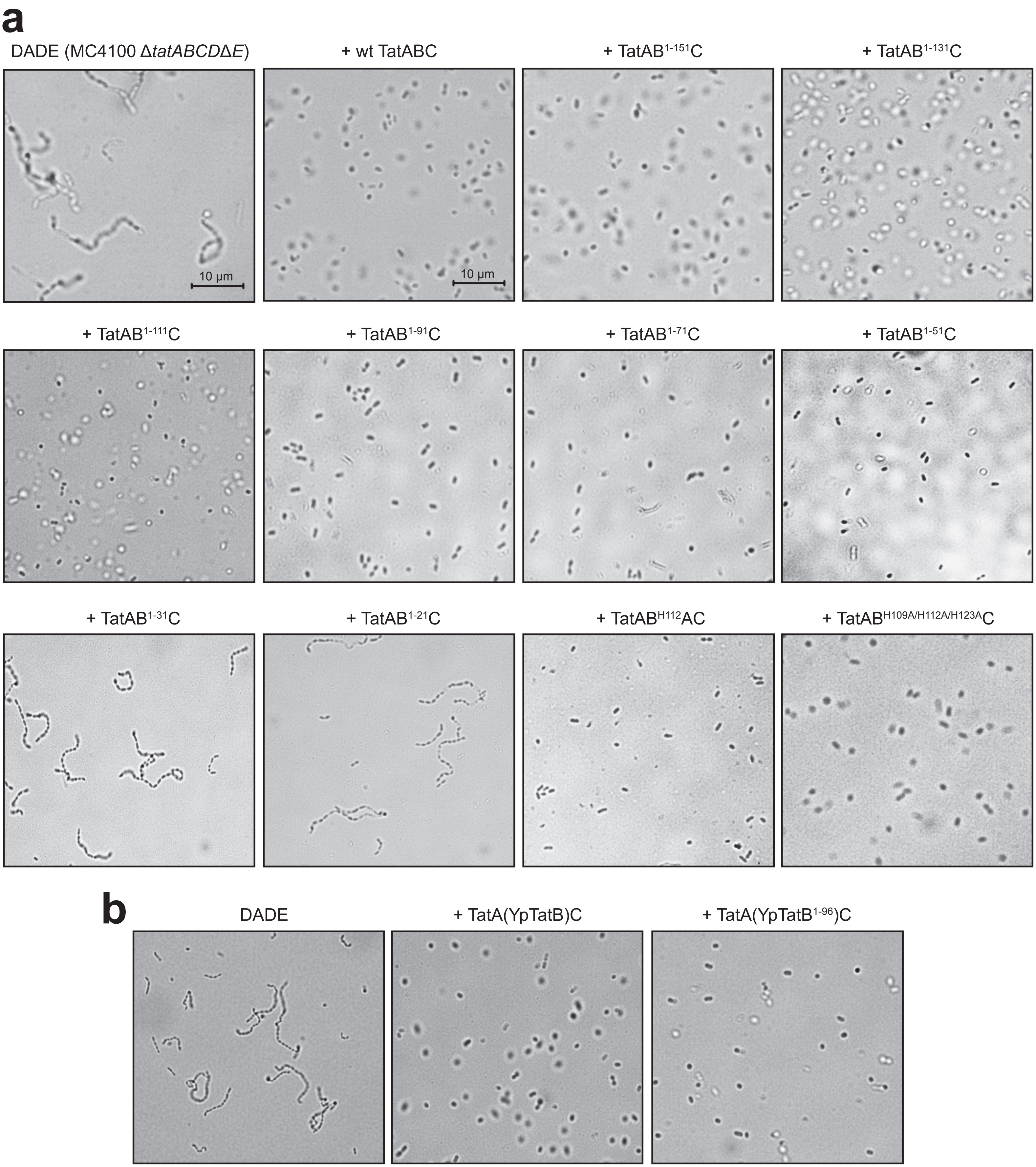


**Supplementary Figure 1. Growth phenotypes conferred by TatB mutants.** (a) Light microscopy of plasmid-free DADE cells or DADE cells expressing wt TatABC or mutant translocases in which TatB was truncated or carrying substitutions in the essential histidine residues as indicated. (b) Light microscopy of plasmid-free DADE cells or DADE cells expressing wt TatAC along with either wt YpTatB or truncated YpTatB$^{1-96}$. Scale bar = 10 µM.

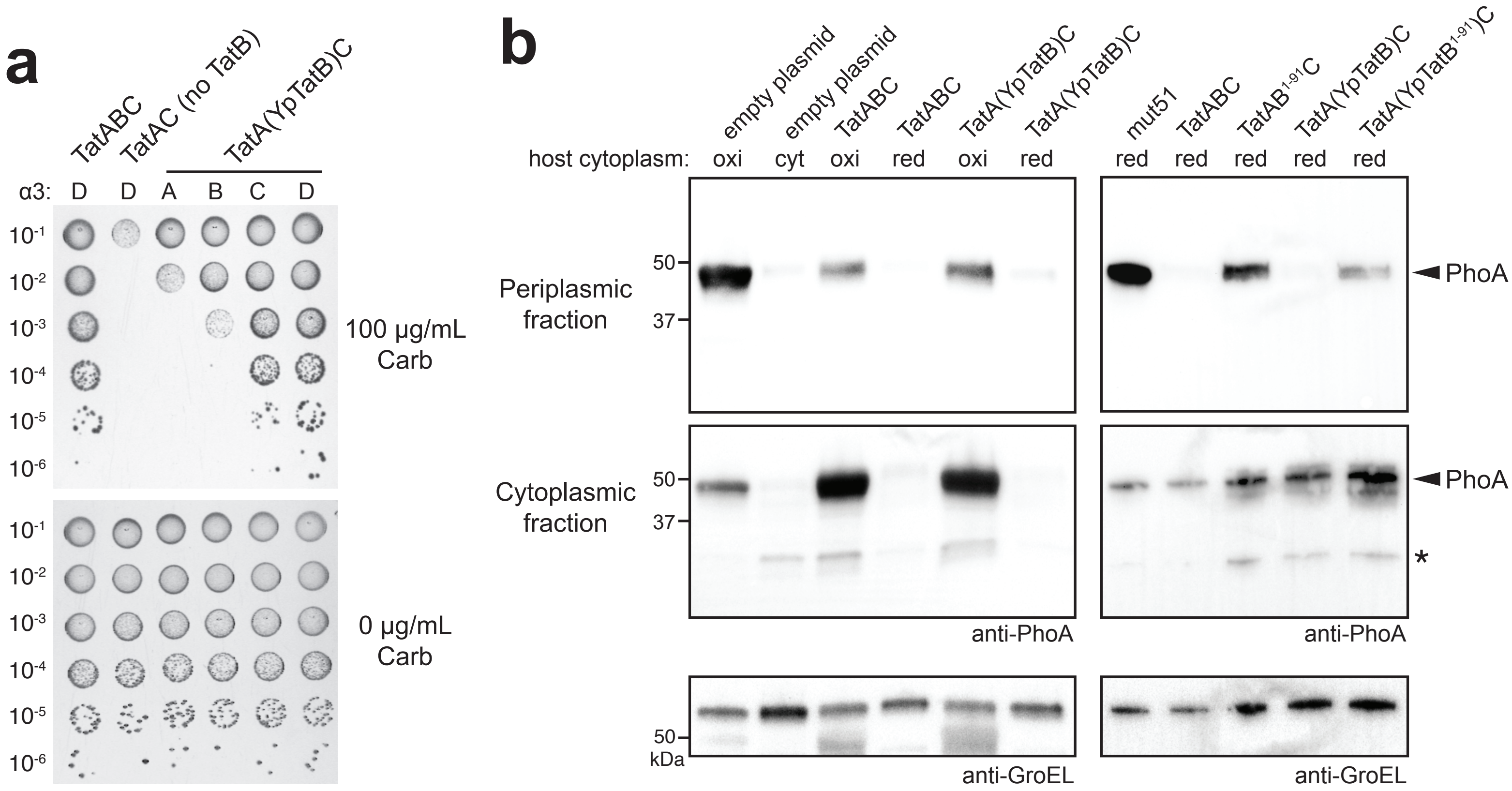


**Supplementary Figure 2. Regulation of QC activity by YpTatB in *E. coli*.** (a) Resistance of serially diluted DADE cells co-expressing one of the spTorA-α3-Bla chimeras (A, B, C or D) along with a Tat operon plasmid encoding wt TatABC or TatAC from *E. coli* without TatB or with YpTatB. Cells were spotted on LB-agar plates containing either 100 µg/mL Amp or 25 µg/mL chloramphenicol (Cam; 0 µg/mL Amp). (b) Western blot analysis of cytoplasmic (cyt) and periplasmic (per) fractions prepared from DRB cells (oxi) or MCMTA cells (red) co-expressing Tat-targeted PhoA from pTorA-AP along with either TatB from plasmid pMAF10-TatB or YpTatB from plasmid pMAF10-YpTatB. DR473 (oxidizing cytoplasm; oxi) and DHB4 (reducing cytoplasm; red) cells carrying empty pMAF10 plasmid served as positive and negative controls, respectively. DADE cells carrying a plasmid encoding QCS suppressor mut51 were also included as a positive control. An equivalent number of cells was loaded in each lane. PhoA was probed with anti-PhoA antibody while anti-GroEL antibody confirmed equivalent loading in each lane.

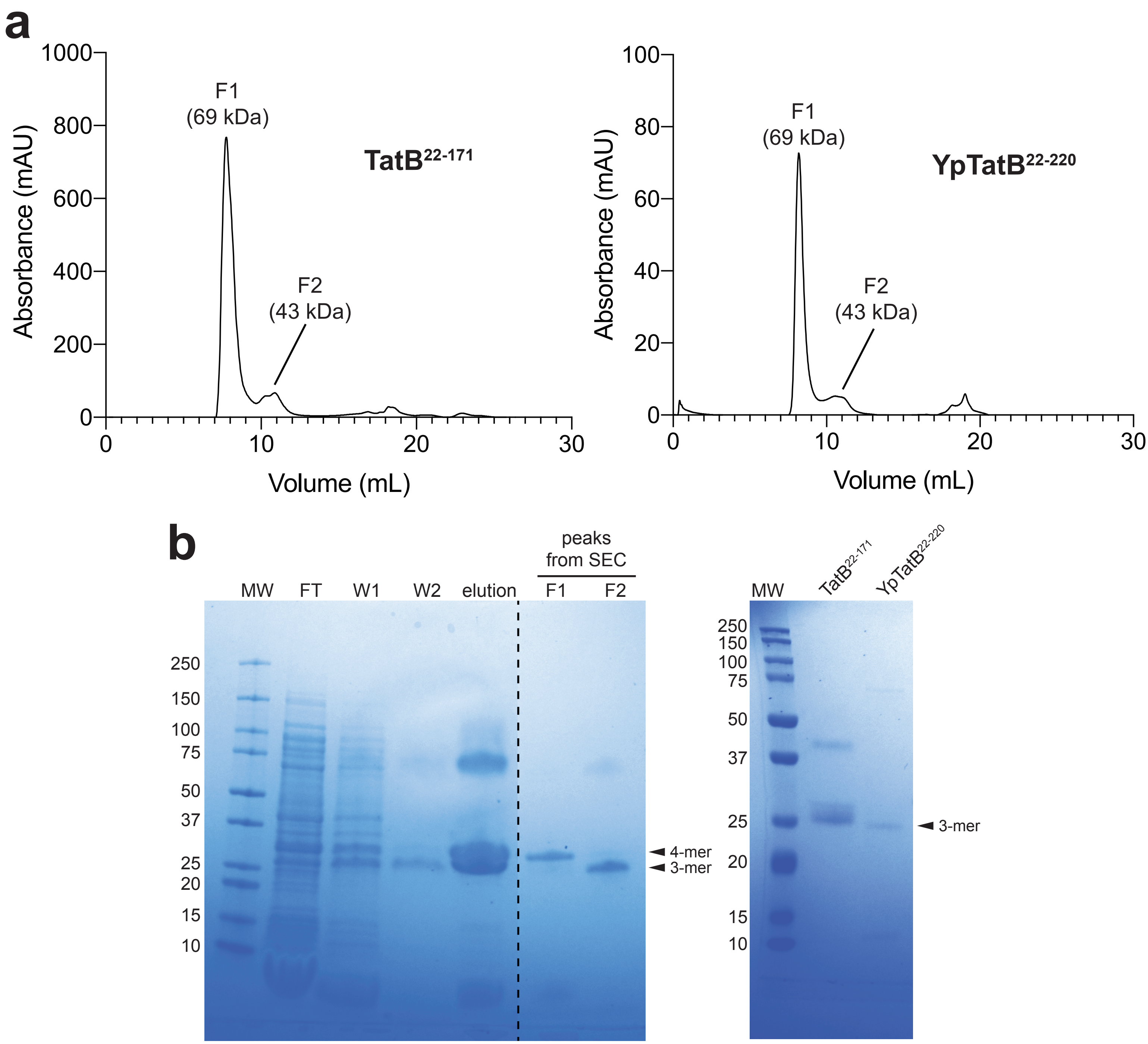


**Supplementary Figure 3. Purification of TatB and YpTatB proteins.** (a) SEC analysis of Ni-NTA-purified *E. coli* TatB$^{22\text{-}171}$ and *Y. pseudotuberculosis* TatB$^{22\text{-}220}$ using GE Superdex 75 10/300 GL. Both purified proteins eluted as oligomers based on the expected molecular weights of TatB$^{22\text{-}171}$ (~17 kDa) and YpTatB$^{22\text{-}220}$ (~23 kDa). (b) Coomassie blue-stained SDS-PAGE gels showing: (left panel) various fractions of TatB$^{22\text{-}171}$ collected during purification along with peak fractions from SEC; and (right panel) final purified TatB$^{22\text{-}171}$ and YpTatB$^{22\text{-}220}$ proteins as indicated.

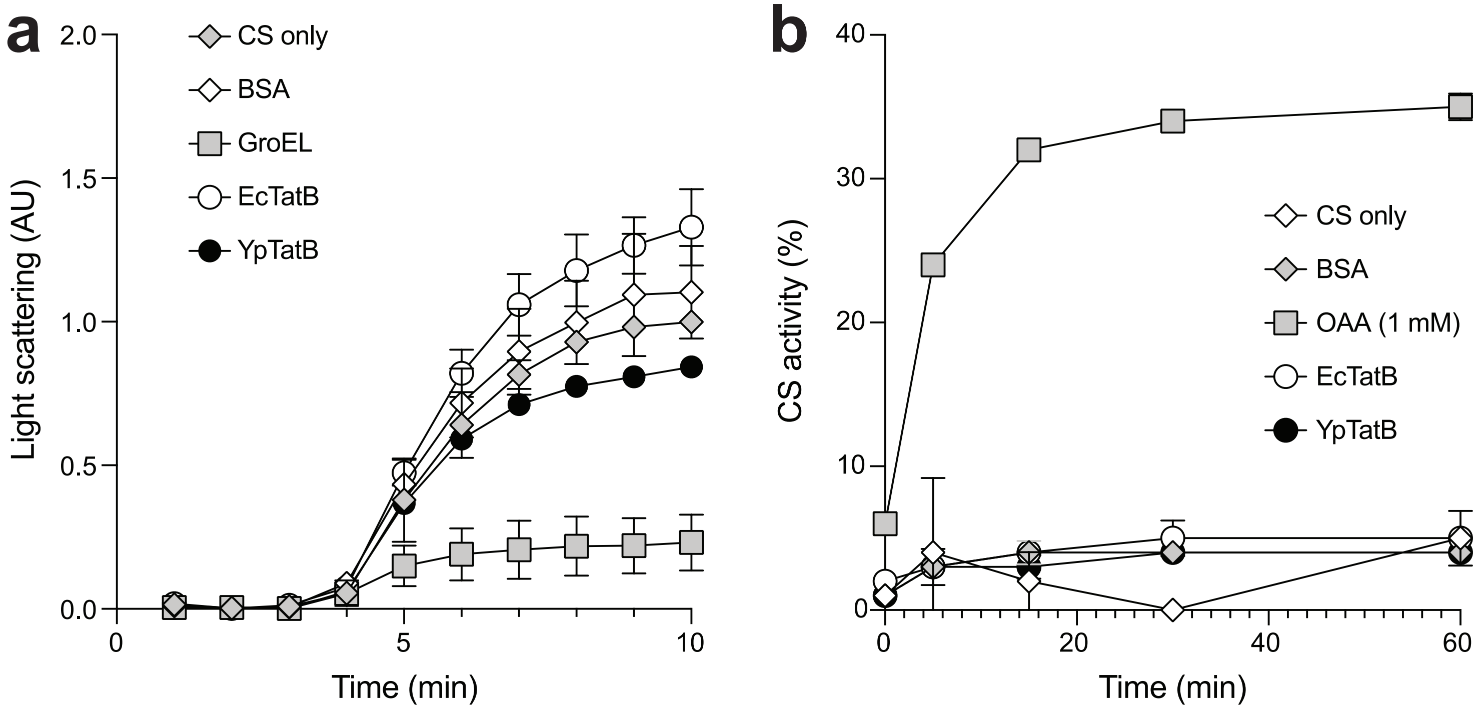


**Supplementary Figure 4. Effect of TatB and YpTatB on aggregation of chemically denatured CS and reactivation of thermally denatured CS.** (a) CS aggregation monitored by light scattering measurements over time without additional factors or in the presence of 0.15 µM of $TatB^{22\text{-}171}$, $YpTatB^{22\text{-}220}$, GroEL, or BSA. CS was chemically denatured in 6 M Gdn-HCl and diluted to a final concentration of 0.15 µM into 50 mM Tris-HCl, pH 8.0. (b) CS (0.15 µM) was inactivated at 43°C in the presence of 0.6 µM $TatB^{22\text{-}171}$, $YpTatB^{22\text{-}220}$, BSA, or 1 mM oxaloacetate (OAA) for 30 min prior to reactivation by transferring samples to room temperature. CS reactivation kinetics were monitored by measuring CS activity at different time points following inactivation. OAA is a substrate of CS and also a known ‘stabilizer’ for slowing down unfolding CS intermediates from irreversible aggregation.

**Supplementary Figure 1. Bacterial strains and plasmids used in this study.**

| **Bacterial Strain** | **Genotype** | **Source** |
|---|---|---|
| MC4100 | F- *araD139* Δ(*argF-lac*)*169* λ- *e14- flhD5301* Δ(*fruK-yeiR*)*725(fruA25) relA1 rpsL150*(Str$^R$) *rbsR22* Δ*(fimB-fimE)632(*::IS*1) deoC1* | (1) |
| DADE | MC4100 Δ*tatABCD*Δ*tatE* | (2) |
| BL21(DE3) | F⁻ *ompT hsdS*$_B$ (r$_B^-$, m$_B^-$) *gal dcm* (DE3) | Laboratory stock |
| MC1000 | F'[*proAB lacI*$^Q$ *lacZ* ΔM15 Tn*10*(Tet$^R$)] *araD139* Δ(*araA-leu*)7679 (*codB-lac*)X74 *galE15 galK16 rpsL150 relA1 thi* | |
| DHB4 | MC1000 Δ*phoA*(*Pvu*II) *phoR* Δ*malF3* | (3) |
| DR473 | DHB4 Δ*trxB gor552* Tn*10*(Tet$^R$) *ahpC** Tn*10*(Cm$^R$) *(araC P$_{ara}$-trxB)* | (4) |
| MCMTA | MC4100 *tatB*::kan | (5) |
| DRB | DR473 *tatB*::kan | (3) |
| **Bacterial plasmid(s)** | | |
| pTatABC | Entire *E. coli tatABC* operon including native promoter in pBR322; Tet$^R$ | (6) |
| pTatABC-XX | pTatABC with XbaI and XhoI restriction sites flanking *tatB* and *tatC* genes, respectively | (6) |
| pTatAC | pBR322 with *tatAC* cloned between PvuI and AhdI | This work |
| pTatAB(-10)C-XX and related TatB truncation plasmids | pTatABC-XX with truncation of *tatB* at 3' end by 10, 20, 30, 40, 50, 60, 70, 80, 90, 100, 110, 120, 130, and 140 codons | This work |
| pTorA-AP | Signal peptide derived from *E. coli* trimethylamine *N*-oxide reductase (TorA) fused to gene encoding mature *E. coli* alkaline phosphatase (APΔ1-22) cloned in plasmid pTrc99A | (3) |
| pET22-TatB$^{22-171}$ | *E. coli tatB* gene lacking N-terminal TMH (residues 1-21) cloned with polyhistidine (6x-His) tag in plasmid pET-22b(+) | This work |
| pET22-YpTatB$^{22-220}$ | *Y. pseudotuberculosis tatB* gene lacking N-terminal TMH (residues 1-21) cloned with polyhistidine (6x-His) tag in plasmid pET-22b(+) | This work |
| pET22-TatB$^{22-91}$ | *E. coli tatB* gene lacking N-terminal TMH and truncated C-terminally to terminate at residue 91 with 6x-His tag in pET-22b(+) | This work |
| pET22-YpTatB$^{22-91}$ | *Y. pseudotuberculosis tatB* gene lacking N-terminal TMH and truncated C-terminally to terminate at residue 91 with 6x-His tag in pET-22b(+) | This work |
| pSALect-α3 plasmids | Codon-optimized genes encoding α3A/B/C/D proteins cloned between spTorA and Bla in plasmid pSALect | (6) |
| pTatA(YpB)C | Gene encoding *Y. pseudotuberculosis* TatB replacing *E. coli* TatB in plasmid pTatABC-XX | This work |
| pTatA(YpB$^{1-91}$)C | Gene encoding C-terminally truncated *Y. pseudotuberculosis* TatB (residues 1-91) replacing *E. coli* TatB in plasmid pTatABC-XX | This work |
| pMAF10-TatB | Gene encoding *E. coli* TatB cloned between EcoRI and XbaI in plasmid pMAF10 | This work |
| pMAF10-YpTatB | Gene encoding *Y. pseudotuberculosis* TatB cloned between EcoRI and XhoI in plasmid pMAF10 | This work |
